# Educational attainment impacts drinking behaviors and risk for alcohol dependence: results from a two-sample Mendelian randomization study with ~780 000 participants

**DOI:** 10.1101/557595

**Authors:** Daniel B. Rosoff, Toni-Kim Clarke, Mark J. Adams, Andrew M. McIntosh, George Davey Smith, Jeesun Jung, Falk W. Lohoff

**Author notes:** Corresponding Author: Falk W. Lohoff, M.D., Chief, Section on Clinical Genomics and Experimental Therapeutics (CGET), Lasker Clinical Research Scholar, National Institute on Alcohol Abuse and Alcoholism (NIAAA), National Institutes of Health, 10 Center Drive (10CRC/2-2352), Bethesda, MD 20892-1540.

## Abstract

Observational studies suggest that lower educational attainment (EA) may be associated with risky alcohol use behaviors; however, these findings may be biased by confounding and reverse causality. We performed two-sample Mendelian randomization (MR) using summary statistics from recent genome-wide association studies (GWAS) with >780 000 participants to assess the causal effects of EA on alcohol use behaviors and alcohol dependence (AD). Fifty-three independent genome-wide significant SNPs previously associated with EA were tested for association with alcohol use behaviors. We show that while genetic instruments associated with increased EA are not associated with total amount of weekly drinks, they are associated with reduced frequency of binge drinking ≥6 drinks (ß_IVW_= −0.198, 95% CI, −0.297-0.099, *P*_IVW_=9.14×10^−5^), reduced total drinks consumed per drinking day (ß_IVW_=−0.207, 95% CI, −0.293-0.120, *P*_IVW_=2.87×10^−6^), as well as lower weekly distilled spirits intake (ß_IVW_=−0.148, 95% CI, −0.188-0.107, *P*_IVW_=6.24×10^−13^). Conversely, genetic instruments for increased EA were associated with increased alcohol intake frequency (ß_IVW_=0.331, 95% CI, 0.267-0.396, *P*_IVW_= 4.62×10^−24^), and increased weekly white wine (ß_IVW_=0.199, 95% CI, 0.159-0.238, *P*_IVW_=7.96×10^−23^) and red wine intake (ß_IVW_=0.204, 95% CI, 0.161-0.248, *P*_IVW_=6.67×10^−20^). Genetic instruments associated with increased EA reduced AD risk: an additional 3.61 years schooling reduced the risk by approximately 50% (OR_IVW_=0.508, 95% CI, 0.315-0.819, *P*_IVW_=5.52×10^−3^). Consistency of results across complementary MR methods accommodating different assumptions about genetic pleiotropy strengthened causal inference. Our findings suggest EA may have important effects on alcohol consumption patterns and may provide potential mechanisms explaining reported associations between EA and adverse health outcomes.

## INTRODUCTION

Alcohol consumption is a major risk factor exhibiting a complex relationship with death and disability in the United States and worldwide with the World Health Organization estimating alcohol is responsible for 139 million disability-adjusted life-years globally^1–3^. Acute intoxication may result in injuries, poisoning, and interpersonal violence^3, 4^, while longer-term alcohol consumption contributes to chronic diseases, including cancer^5–7^, cardiovascular disease^8–10^, and dependent drinking exacerbating psychiatric comorbidities or other impairments^2, 11^. The complex relationship between alcohol and morbidity is due, in part, to the pattern of its use and the beverage type consumed^12, 13^ with beer and hard liquor consumption associated with more severe drinking patterns and increased risk for alcohol-related problems^14^. In addition, while the quantity of alcohol consumed and alcohol intake frequency are correlated ^15^, and often used interchangeably, they demonstrate different and often opposite effects on health^15^. Using genetic correlations, Marees et al. 2019 recently showed opposing associations of alcohol quantity and intake frequency with many health behaviors, including smoking, various psychiatric disorders, and personality traits suggesting different risk profiles related to these alcohol consumption metrics^15^.

The suggested differences in risk profiles with different alcohol consumption patterns and the seriousness of the acute and chronic diseases linked with risky alcohol consumption highlights the importance of identifying causal risk factors related to how alcohol is consumed to develop and improve intervention and treatment strategies. Among various social determinants associated with health disparities (age, gender, race, ethnicity, etc.) and mortality, educational attainment (EA) has been identified as a prominent risk factor^16^. For example, at age 25, the average life expectancy of U.S. adults without a high school diploma is nine years shorter than college graduates^17^. The impact of education on alcohol consumption behaviors may be an important pathway mediating these effects. Observational studies have demonstrated EA likely influences drinking patterns, beverage preferences, and alcohol-related outcomes^13, 18, 19^. Higher EA is associated with reduced odds of reporting high-risk drinking^1, 18^, or at least one episode of heavy episodic drinking within the past twelve months^13, 17, 20^. Moreover, individuals with fewer years of education are more likely to report higher single occasion quantity consumed and alcohol-related harm^17^. There is also conflicting evidence that alcohol consumption associates with EA with some studies showing an association with decreased years of schooling while others find either very small or non-significant effects^21^. While education is associated with differences in alcohol consumption behaviors, observational studies are subject to reverse causation, or residual confounding^22–24^. Recent genetic studies have suggested that the relationship between alcohol use and EA is complex^15^ and differs markedly depending on which aspect of alcohol use is considered. Sanchez-Roige et al. (2019) observed a positive genetic correlation (r_g_) between college completion and Alcohol Use Disorder Identification Test (AUDIT)^25^ total scores (r_g_=0.23; standard error (se)=0.05)^26^; Walters et al. (2016) observed a negative genetic correlation between EA and DSM-IV alcohol dependence (AD) (r_g_= −0.47; se=0.07)^27^; and Marees et al. (2019) found a positive genetic correlation between EA and alcohol intake frequency^26, 28^ but a negative correlation between EA and total alcohol intake quantity^27^. (Genetic correlations for the current study are presented in Supplementary Table 1.) Furthermore, inferring causality from correlations and multivariable adjusted regression models is often unreliable^29, 30^.

Mendelian randomization (MR) analysis uses randomly inherited genetic markers (single nucleotide polymorphisms (SNPs)) robustly associated with a risk factor (*e.g.*, education) as proxies for environmental exposures to assess causal inferences about the effect of the exposure on an outcome (*e.g.*, alcohol consumption patterns and AD risk). MR has some analogies to randomized controlled trials (RCTs) since genetic variants are not modifiable and free from reverse confounding^23, 24, 31^ and is an important strategy for establishing evidence of causal relationships where RCTs are impractical or unethical^23^. The increasing availability of summary-level data from genome-wide association studies (GWASs) can be used to perform MR analyses where gene exposure and gene outcome measures are derived from two separate GWAS^32^. These two-sample MRs benefit from increased statistical power and enable sensitivity analyses to test the robustness of the MR findings^33^.

Recent two-sample MR studies have shown inverse relationships between EA and both smoking and coronary heart disease^34, 35^. However, to our knowledge, MR has not been applied to examine the effects of EA on alcohol consumption patterns, DSM-IV alcohol dependence, or other indicia of alcohol use disorders. In this study, using the largest, publicly available GWASs to date, we conduct a bidirectional two-sample MR of EA (N=293 723) ^36^ on AD (N=28 657)^27^, AUDIT scores^25^ (N ≤ 121 604)^26^, and alcohol consumption (total quantity consumed^37^, intake frequency, whether alcohol is consumed with meals, and drink-specific average weekly intake (cider and beer, red and white wine, and distilled spirits)) (N ≤ 462 346)^38^ to assess evidence of causal associations between EA and alcohol dependence and consumption. Given men and women differ in their alcohol consumption patterns and alcohol-related problems^39^, we also performed exploratory two-sample MR analyses using sex-specific alcohol consumption and AUDIT GWASs (N (females) ≤ 194 174; N (males) ≤ 167 010), where available, to evaluate whether EA differentially impacts drinking behaviors between men and women.

## METHODS

### DATA SOURCES

GWASs included in the current study are described in Table 1. We selected online publicly available GWASs with the largest sample sizes consisting of populations of European ancestry and without significant sample overlap. Details of the GWASs, including quality control and association methods, are available in Supplementary Methods 1. All GWASs have existing ethical permissions from their respective institutional review boards and include participant informed consent.

**Table 1.** GWASs included in the current study.

| **PHENOTYPE** | **Source** | **Citation** | **Sample Size** | **Variable** |
| --- | --- | --- | --- | --- |
| **Educational attainment (SD=3.61years)** | SSGAC | Okbay et al. (2016) | 293 723 | Continuous |
| **Average before tax household income** | Neale Lab UKB | www.nealelab.is/uk-biobank | 411 028 | Categorical |
| **ALCOHOL USE:** |  |  |  |  |
| **Alcohol intake frequency** | MRC-IEU UKB | Mitchell et al. (2019) | 462 346 | Categorical |
| **Weekly intake (drinks per week)** | SSGAC | Karlsson Linnér (2019) | 414 343 | Integer |
| **Weekly intake by drink type (units):** |  |  |  |  |
| **Distilled spirits (measure)** | MRC-IEU UKB | Mitchell et al. (2019) | 326 565 | Categorical |
| **Beer plus cider (pint)** | MRC-IEU UKB | Mitchell et al. (2019) | 327 634 | Categorical |
| **Red wine (glass)** | MRC-IEU UKB | Mitchell et al. (2019) | 327 026 | Categorical |
| **Champagne plus white wine (glass)** | MRC-IEU UKB | Mitchell et al. (2019) | 326 801 | Categorical |
| **ALCOHOL DEPENDENCE (AD):** |  |  |  |  |
| **Alcohol dependence (DSM-IV diagnosis)** | PGC | Walters et al. (2018) | 28 657 | Binary |
| **ALCOHOL USE DISORDERS IDENTIFICATION TEST (AUDIT):** |  | www.nealelab.is/uk-biobank |  |  |
| **Frequency of alcohol intake** | Neale Lab UKB |  | 117 914 | Categorical |
| **Amount of alcohol drunk on a typical drinking day** | Neale Lab UKB |  | 108 256 | Categorical |
| **Frequency of consuming $\geq 6$ or more units of alcohol** | Neale Lab UKB |  | 108 485 | Categorical |
| **Frequency of inability to cease drinking in the last year** | Neale Lab UKB |  | 67 973 | Categorical |
| **Frequency of failure to fulfill normal expectations due to alcohol (past year)** | Neale Lab UKB |  | 65 054 | Categorical |
| **Frequency of needing a morning drink** | Neale Lab UKB |  | 65 099 | Categorical |
| **Frequency of feeling guilt or remorse after drinking alcohol (past year)** | Neale Lab UKB |  | 65 009 | Categorical |
| **Frequency of memory loss due to drinking alcohol (past year)** | Neale Lab UKB |  | 65 029 | Categorical |
| **Ever been injured or injured someone else** | Neale Lab UKB |  | 118 002 | Categorical |
| **Ever had a known person concerned or recommend reduction** | Neale Lab UKB |  | 117 880 | Categorical |

### GENETIC INSTRUMENTS FOR EXPOSURE: EDUCATIONAL ATTAINMENT

We extracted summary association statistics for the 74 genome-wide significant (*P*<5×10^−8^) SNPs previously demonstrated to be associated with EA defined as years of schooling ^36^, measured in standard deviation (sd) units (mean=14.33 years, sd=3.61 years), ascertained in up to 293 723 persons from 64 discovery cohorts (excluding the subsequent UKB replication cohort). Twenty-one of the 74 SNPs were excluded by linkage disequilibrium (LD) of R^2^=.001 and clumping distance=10 000 kb, leaving 53 independent SNPs for the two-sample MR analysis (Supplementary Table 2).

EA is highly heritable and strongly genetically correlated with markers of socioeconomic status (SES)^40^. Non-zero genetic correlations suggest shared genetic factors contributing to these social outcomes^40^. We sought to account for the effects of SES, as operationalized by income, by performing additional analyses with an EA instrument constructed by removing variants associated with income. We took advantage of the recent average household before tax income GWAS (N=311 028) from the Neale Lab UK Biobank (UKB) GWAS (http://www.nealelab.is/uk-biobank/), generating an ordinal categorical phenotype (sample frequency in %): < 18 000£ (21.8%) , 18 000–30 999£ (25.4%), 31 000–51 999£ (26.4%), 52 000–100 000£ (21.9%), and > 100 000£ (5.5%). We removed variants from the EA instrument associated with income at a threshold significance of .00094 (nominal *P*=.05 corrected for 53 comparisons, the number of SNPs in the main EA instrument), leaving 30 independent variants (Supplementary Table 3).

### GENETIC INSTRUMENTS FOR OUTCOMES

#### Alcohol consumption

We used summary statistics from the MRC-IEU UKB GWAS Pipeline^38^ on six alcohol consumption behaviors generated using the PHEnome Scan Analysis Tool (PHESANT)^41^ as ordinal categorical responses: (1) alcohol intake frequency, *i.e.* never, special occasions only, one to three times a month, once or twice a week, three or four times a week, daily or almost daily (N=432 346); and for the subset of UKB respondents who indicated they drank at least once or twice a week, also assessed were (2) average weekly spirits intake in increments of one pub measure of alcohol (N=326 565); (3) average weekly beer plus cider intake in increments of one pint (N=327 634); (4) average weekly red wine intake in increments of one glass (N=327 026); (5) average weekly white wine plus champagne in increments of one glass (N=326 801); and (6) average weekly fortified wine intake in increments of one glass (N=327 563). We also used summary association statistics from the MRC-IEU UKB GWAS Pipeline on an additional alcohol consumption behavior assessed as a binary response (0=No, 1=Yes): (7) alcohol usually taken with meals (N cases=159 104; N controls=75 541). For the additional sex-specific analyses, we used statistics from sex-specific GWASs from the Neale Lab UKB GWAS: sex-specific GWASs were not available from the MRC-IEU UKB GWAS Pipeline. Out of the 53 possible independent SNPs associated with EA, 52 were present in these MRC-IEU GWASs, 1 SNP was identified in high linkage disequilibrium (LD) as a proxy for the missing SNP, and 2 SNPs were removed for being palindromic with intermediate allele frequencies (to harmonize the data so that the effect of the variants on the exposure EA and outcomes corresponded to the same allele), leaving 51 SNPs for analysis (Supplementary Table 2).

For a measure of total alcohol consumption, we used summary statistics from the SSGAC GWAS of alcohol consumption in the UKB (N=414 343), measured as “drinks per week” (DPW), constructed, for UKB participants who indicated they drank “at least once or twice per week”, by aggregating the weekly intake of distilled spirits (pub measures), beer and cider (pints), red wine, white wine, and champagne (glasses), and other alcoholic drinks *e.g*. alcopops (DPW: mean=8.92 drinks, SD=9.30 drinks) ^37^. For UKB participants who indicated they drank “one to three times a month”, the phenotype was constructed by aggregating the monthly intake over all drink types and dividing by four. Sex-specific alcohol consumption GWASs were not available from the SSGAC. Out of the 53 possible SNPs associated with EA, 52 were present in this SSGAC GWAS, and 2 SNPs were removed for being palindromic with intermediate allele frequencies, leaving 50 SNPs (Supplementary Table 2).

#### Alcohol dependence and alcohol use disorders identifiers

We used summary association statistics from the PGC GWAS on AD, defined as meeting criteria for a DSM-IV (or DSM-IIIR in one instance) diagnoses, in 28 657 participants (cases N=8 485; controls N=20 657)^27^. Sex-specific AD GWASs were not available. Out of the 53 possible SNPs associated with EA, 53 were present in this PGC GWAS, and 9 SNPs were removed for being palindromic with intermediate allele frequencies, leaving 44 SNPs (Supplementary Table 2).

To assess identifiers or symptoms of alcohol dependence or use, we used summary association statistics from the Neale Lab UKB GWAS (http://www.nealelab.is/uk-biobank/) for responses to the ten-item AUDIT; for the supplementary sex-specific analyses, we used statistics from the corresponding sex-specific GWASs. The PHESANT^41^ generated phenotype categories are further described in Supplementary Table 4 and Supplementary Methods 2. Out of the 53 possible SNPs associated with EA, 53 were present in these ten Neale Lab GWASs, and 8 SNPs were removed for incompatible alleles, leaving 45 SNPs (Supplementary Table 2).

### BIDIRECTIONAL ANALYSIS

We extracted exposure summary association statistics for genome-wide significant (*P* < 5×10^−8^) SNPs associated with the alcohol consumption, AD and AUDIT GWASs described above, removed SNPs in LD with other SNPs, then extracted outcomes in the EA GWAS, harmonized exposure and outcomes, removing palindromic alleles with intermediate frequencies, for the bidirectional two-sample MR analyses. Further details are described in Supplementary Methods 3.

### SAMPLE OVERLAP

Participant overlap between the samples used to estimate genetic associations between exposure and outcome in two sample MR can bias results^42^. We endeavored to use only non-overlapping GWAS summary statistics to reduce this source of bias. We used the discovery EA GWAS (N=293 723) including the 23&Me cohort but excluding the UKB replication cohort; thus, there was no overlap between the exposure cohorts and alcohol consumption and AUDIT outcomes (solely UKB cohorts). For alcohol dependence, a comparison of the cohorts included in the PGC alcohol dependence GWAS and SSGAC EA GWAS showed 2 common cohorts (N=11 096) (Minnesota Center for Twin and Family Research; Swedish Twin Registry). The relevant percentage overlap for purposes of determining weak instrument bias (WIB) is taken with respect to the larger data set – only the presence of participants in both studies leads to correlation in estimates^42^. Here, the two common cohorts accounted for 3.8% of participants in the larger SSGAC GWAS; based on simulation studies of the association between sample overlap and the degree of WIB, considerable bias is not expected^42^.

### STATISTICAL ANALYSIS

#### Genetic correlation

We estimated SNP heritability as well as cross-trait genetic correlation between EA and alcohol consumption and dependence by linkage disequilibrium score regression (LDSR)^43^, using summary level statistics from the previously conducted GWASs, all based on large samples. We used the centralized database and web interface, LD Hub^44^. Analysis was restricted to well-imputed SNPs for the selected phenotypes, with SNPs filtered to HapMap3 SNPs with 1000 Genomes EUR MAP above 5%, and insertions and deletions, structural variants, strand-ambiguous and unmatched SNPs removed, along with SNPs within the major histocompatibility complex region, and SNPs with extremely large effect sizes^44^. Significant genetic correlations within the UKB cohort were identified by applying a Bonferroni correction for 20 cross-trait comparisons (threshold *P* <.0025) (Supplementary Table 1).

#### Two-sample Mendelian randomization

We used four complementary methods – inverse variance-weighted (IVW) MR, MR Egger, weighted median, and weighted mode MR–to assess evidence of the association of EA and the risks of alcohol use behaviors and alcohol use disorders and also discern sensitivity to different patterns of violations of instrumental variable (IV) assumptions ^33^. We reference IVW MR for the main results: in the absence of pleiotropy and assuming the instruments are valid, IVW MR estimates are the best unbiased estimates ^45^. Consistency of results across these methods (each making different assumptions about pleiotropy) strengthens causal inference; significant divergent results may indicate bias from genetic pleiotropy.

To evaluate heterogeneity in instrument effects, which may indicate potential violations of the IV assumptions underlying two-sample MR^46^, we used both MR Egger intercept test^46^ and the Cochran heterogeneity test^47^. We also used the MR pleiotropy residual sum and <u>o</u>utlier (MR-PRESSO) global test^48^ to identify outlier variants for removal to correct potential directional horizontal pleiotropy and resolve detected heterogeneity. We include an overview of the analyses in Figure 1. Details about these MR methods and tests are included in Supplementary Methods 4. We used the Steiger directionality test to test the causal direction between the hypothesized exposure and outcomes^49^. Given a nominal threshold of .05, and 20 comparisons in the UKB sample, we apply Bonferroni corrected threshold *P*=.0025. Analyses were carried out using TwoSampleMR, version 4.16^33^, and MR-PRESSO, version 1.0^50^, in the R environment, version 3.5.1 (2018-07-02).

**Figure 1.**
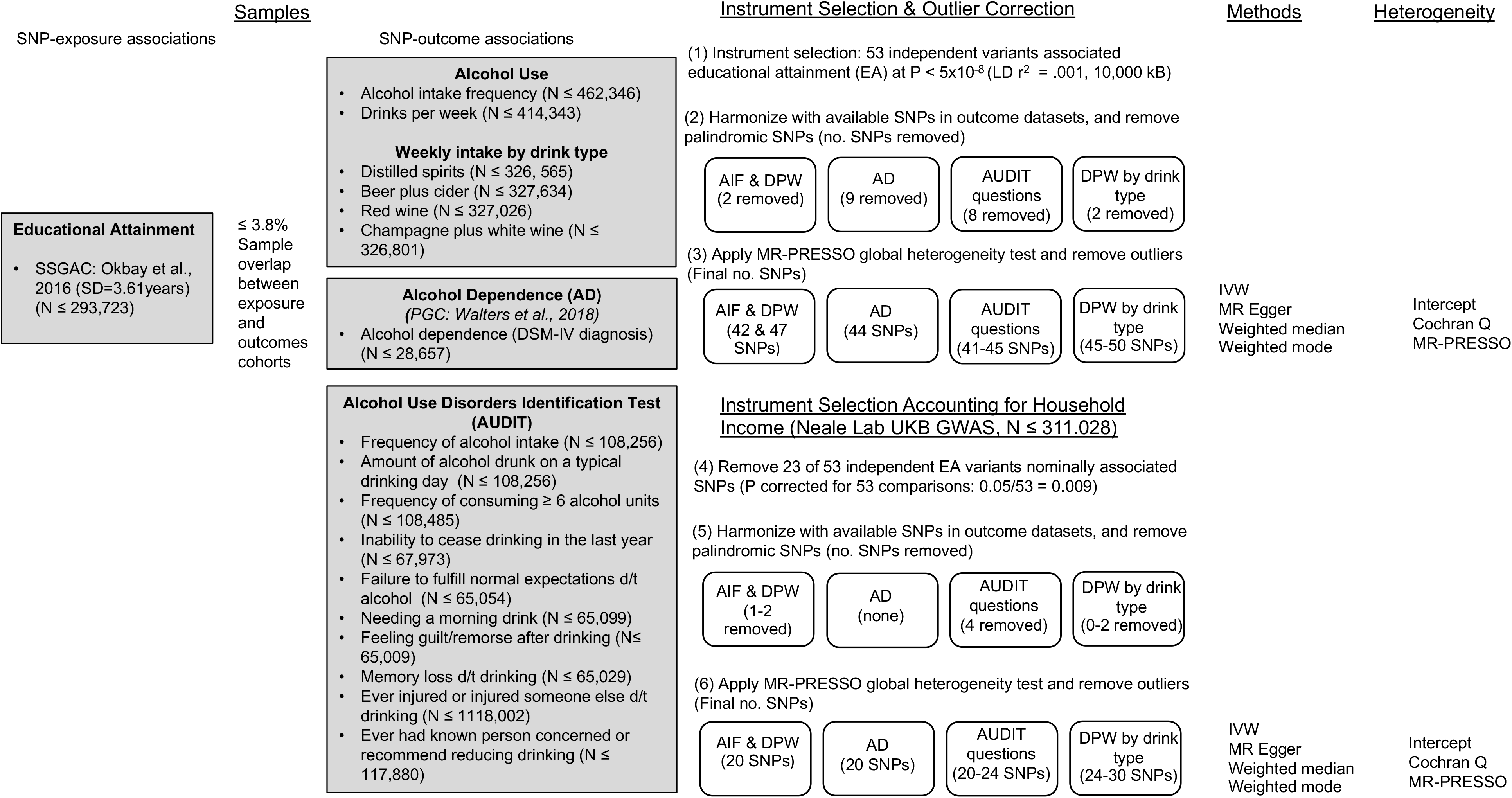
Overview of the main analysis. Abbreviations: SSGAC=Social Science Genetics Association Consortium; PGC=Psychiatric Genomics Consortium; MRC-IEU=Medical Research Council Integrative Epidemiological Unit, University of Bristol; GWAS=genome wide association study; UKB=UK Biobank; AIF= alcohol intake frequency; DPW=drinks per week; AD=alcohol dependence; AUDIT=Alcohol Use Disorder Inventory Test; SNP = single nucleotide polymorphism; N = sample size; MR = Mendelian randomization; IVW = inverse weighted variance Mendelian randomization.

## RESULTS

### OVERVIEW

We present the genetic correlation results from LDSR in Supplementary Table 1. As regards two-sample MR, we report those estimates (1) agreeing in direction and magnitude across MR methods, exceeding nominal significance (*P* <.01) in IVW MR, (2) not indicating bias from horizontal pleiotropy (MR-PRESSO global test *P* >.01), nor directional pleiotropy (MR Egger intercept *P* >.01), and (3) indicating true causal effect directionality (Steiger directionality test *P* <.01), except where otherwise noted. We present the outlier corrected MR estimates in Figures 2 and 3; individual genetic variant associations in Supplementary Table 2; MR results in Supplementary Tables 5-7; bidirectional MR results in Supplementary Table 8; and in Supplementary Table 9, single-SNP and leave-one-out results for the main analyses.

**Figure 2.**
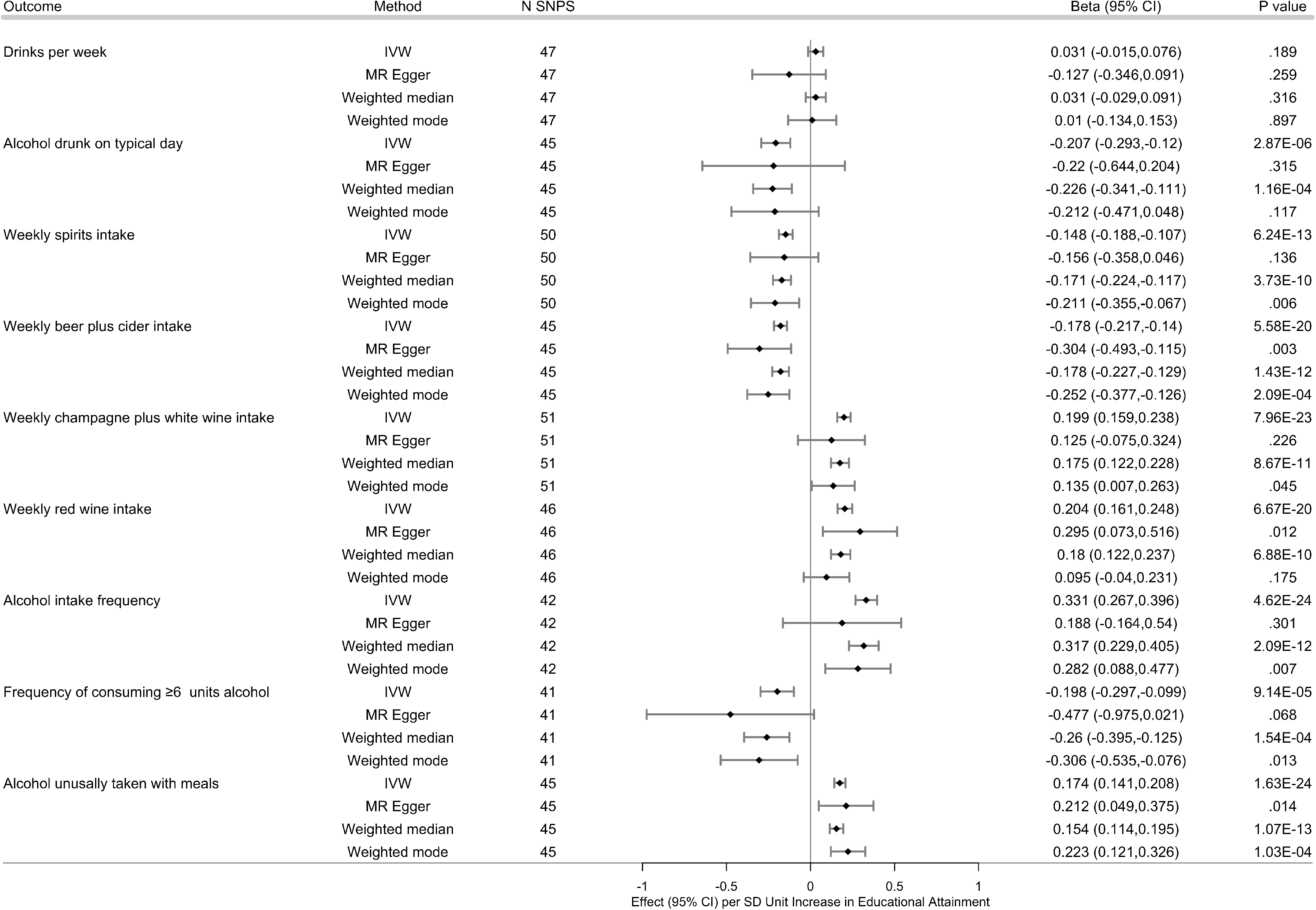
Effects of the genetic variants for increased educational attainment (EA) on alcohol use. 53 genome-wide significantly associated (*P* < 5×10^−8^) independent (LD R^2^ = .001, clumping distance=10 000 kb) single nucleotide polymorphisms (SNPs) were used as instruments for EA. Results from inverse variance-weighted (IVW) and three complementary two-sample MR methods, following removal of variants identified as outliers (MR PRESSO *P* < .10), are shown. Effect (ß) measures the change per unit increase in outcome per standard deviation (SD=3.61 years) increase in EA. Error bars indicate 95% confidence intervals at the nominal threshold .05. With 20 comparisons overall in the UKB cohort, the Bonferroni corrected threshold for comparisons would be .0025, given a nominal threshold .05. Abbreviations: LD=linkage disequilibrium; MR= Mendelian randomization; IVW=Inverse Variance Weighted MR; ß=effect estimate.

**Figure 3.**
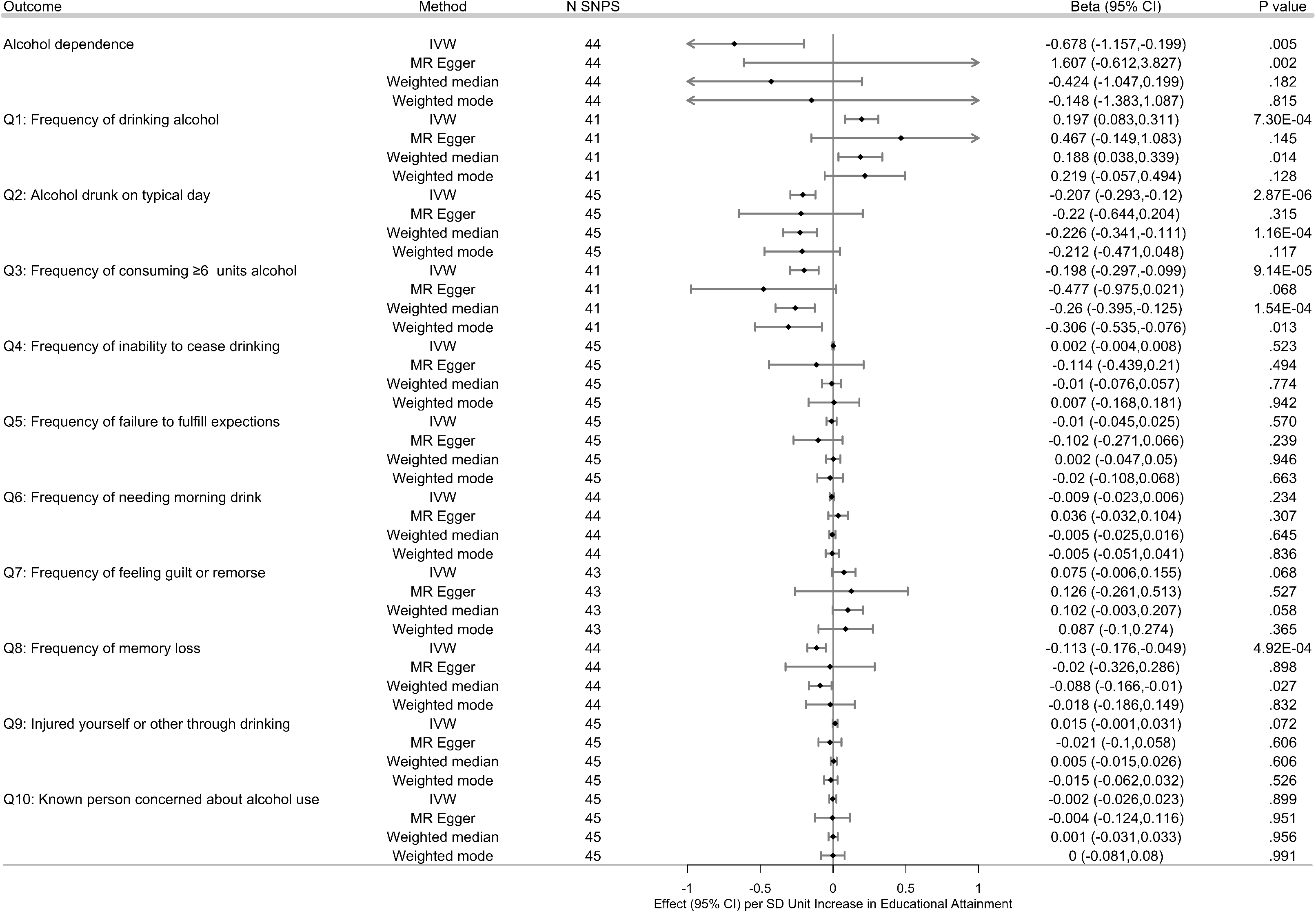
Effects of the genetic variants for increased educational attainment (EA) on alcohol dependence (AD) and AUDIT. 53 genome-wide significantly associated (*P* < 5×10^−8^) independent (LD R^2^ = .001, clumping distance=10 000 kb) single nucleotide polymorphisms (SNPs) were used as instruments for EA. AUDIT outcomes were assessed on sub-cohort of UKB participants in the UKB AUDIT module. Results from inverse variance-weighted (IVW) and three complementary two-sample MR methods, following removal of variants identified as outliers (MR PRESSO *P* < .10) are shown. Effect (ß) measures the change per unit increase in outcome per standard deviation (SD=3.61 years) increase sin EA; with regard to binary outcome AD, ß would be equal to the ln (OR) of AD per SD unit increase in EA. Error bars indicate 95% confidence intervals at the nominal threshold .05. With 20 comparisons overall in the UKB cohort, the Bonferroni corrected threshold for comparisons would be .0025, given a nominal threshold .05. With only one comparison in the PGC AD study, the threshold for AD is .05. Abbreviations: EA=Educational Attainment; AUDIT = Alcohol Use Disorder Identification Test: MR= Mendelian randomization; IVW=Inverse Variance Weighted MR; ß=effect estimate; LD=linkage disequilibrium; OR=odds ratio.

### GENETIC CORRELATIONS FROM LDSR

EA showed significant genetic correlations (Bonferroni corrected *P* <.0025 for 20 comparisons within the UKB cohort) with all of the alcohol intake (quantity) traits: positive weak correlation with total drinks per week (r_g_=.092, *P*=4.75×10^−5^), but negative weak correlation with amount of alcohol drunk on a typical drinking day (AUDIT question 2: r_g_=−.240, *P*=1.89×10^−10^); as well as correlations, in different directions, of EA with alcohol intake by drink type: positive strong correlations with average weekly intake of champagne plus white wine (r_g_=.654, *P*=5.86×10^−109^) and red wine (r_g_=.609, *P*=9.99×10^−154^), negative moderate correlations with average weekly intake of distilled spirits (r_g_=−.372, *P*=8.06×10^−32^) and beer plus cider (r_g_=−.409, *P*=2.79×10^−40^). We also observed significant positive genetic correlations of EA with a subset of alcohol intake frequency traits: moderate correlation with alcohol intake frequency (r_g_=.465, *P*=1.99×10^−133^), strong correlation with alcohol usually taken with meals (r_g_=.622, *P*=1.67×10^−186^), but none with frequency of consuming ≥6 alcohol units.

EA also showed significant negative moderate genetic correlation with the AD risk (r_g_=−.463, *P*=6.82×10^−10^); but weak correlations, in opposite directions, with two of four AUDIT responses related to problematic or hazardous alcohol use (frequency of feeling guilt or remorse after drinking (past year): r_g_=.164, *P*=.0007; frequency of memory loss due to drinking alcohol (past year): r_g_=−.203, *P*=.0003). We did not observe evidence for a significant genetic correlation of EA and the three AUDIT responses related to AD. We did observe a strong positive correlation of EA with average household income before tax (r_g_=.805, *P* <.001). See Supplementary Table 1.

### EFFECTS OF EDUCATIONAL ATTAINMENT ON ALCOHOL CONSUMPTION AND CONSUMPTION FREQUENCY

#### Weekly alcohol intake

Genetic variants associated with increased EA were not significantly associated with total number of drinks per week (sum total of different types of drinks) (ß_IVW_=0.031, 95% CI, −0.015-0.076, *P*_IVW_=.189). In the subsample of UKB participating in the AUDIT module, variants associated with increased EA, however, were associated with decreased alcohol intake on a typical drinking day (AUDIT question 2: ß_IVW_=−0.207, 95% CI, −0.293- -0.120, *P*_IVW_=2.87×10^−6^), Disaggregating DPW by drink type, variants associated with increased EA were also associated with decreased average weekly spirits intake (ß_IVW_=−0.148, 95% CI, −0.188- -0.107, *P*_IVW_=6.24×10^−13^) and weekly beer plus cider intake (ß_IVW_=−0.178, 95% CI, −0.217- -0.140, *P*_IVW_=5.58×10^−20^), although outlier correction notwithstanding, evidence of residual heterogeneity and horizontal pleiotropy for beer plus cider intake did remain. Conversely, variants associated with increased EA were associated with increased average weekly champagne plus white wine intake (ß_IVW_=0.199, 95% CI, 0.159-0.238, *P*_IVW_=7.96×10^−23^) as well as increased average weekly fortified wine (ß_IVW_=0.050, 95% CI, 0.027-0.073, *P*_IVW_=1.87×10^−5^); and increased average weekly red wine intake (ß_IVW_=0.204, 95% CI, 0.161-0.248, *P*_IVW_=6.67×10^−20^). See Supplementary Table 5.

#### Alcohol intake frequency

Genetic variants associated with increased EA were associated with increased alcohol intake frequency (ß_IVW=_0.331, 95% CI, 0.267-0.396, *P*_IVW_= 4.62×10^−24^; *see also* AUDIT question 1: ß_IVW_=0.197, 95% CI, 0.083-0.311, *P*_IVW_= 7.30×10^−4^). Variants associated with increased EA were also associated with decreased frequency of consuming six or more units of alcohol per occasion (AUDIT question 3: ß_IVW_=−0.198, 95% CI, −0.297-0.099, *P*_IVW_=9.14×10^−5^), and also with an increased probability of drinking alcohol with meals (ß_IVW_=0.174, 95% CI, 0.141-0.208, *P*_IVW_=1.63×10^−24^), although evidence of residual heterogeneity and horizontal pleiotropy did remain (Supplementary Table 5).

#### Alcohol intake and intake frequency accounting for income

Removing EA instruments associated with average household income attenuated the significance, and reduced heterogeneity, but with only one exception, did not significantly affect the magnitude nor direction of the associations. Variants associated with increased EA, but not associated with household income, were associated with increased average weekly red wine intake, but with a smaller effect size (ß_IVW_=0.123, 95% CI, 0.064-0.181, *P*_IVW_=3.80×10^−5^) (Supplementary Table 6A).

#### Sex-specific intake and intake frequency

Exploratory sex-specific analyses were motivated by these results, *i.e.* EA differentially associated with average weekly intake by drink types, along with surveys finding drink of choice differs by sex, females preferring wine, then beer and spirits, and males preferring beer, then spirits, lastly wine ^51^. Genetic variants associated with increased EA were differentially associated across sexes (non-overlapping CIs) with decreased average weekly spirits intake, with greater effect for females (female: ß_IVW_=−0.218, 95% CI, −0.286--0.150, *P*_IVW_=2.95×10^−10^; male: ß_IVW_=−0.084, 95% CI, −0.146--0.022, *P*_IVW_=7.65×10^−3^); and also decreased average weekly beer plus cider intake, with, in contrast, greater effect for males (female: ß_IVW_=−0.115, 95% CI, −0.165--0.065, *P*_IVW_=5.91×10^−6^; male: ß_IVW_=− 0.246, 95% CI, −0.321--0.170, *P*_IVW_=1.75×10^−10^); but with increased average weekly intake of red wine, again with greater effect for males (female: ß_IVW_=0.161, 95% CI, 0.094-0.228, *P*_IVW_=2.61×10^−6^; male: ß_IVW_=0.262, 95% CI, 0.200-0.324, *P*_IVW_=1.45×10^−16^). Variants associated with increased EA were associated with average weekly intake of white wine plus champagne for both females and males, with no significant difference in effect between sexes (overlapping CIs) (female: ß_IVW_=0.195, 95% CI, 0.129-0.261, *P*_IVW_=5.97×10^−9^; male: ß_IVW_=0.205, 95% CI, 0.144-0.266, *P*_IVW_=5.16×10^−11^). See Supplementary 6B. Sex-specific GWASs for drinks per week were not available; however, in the subsample of UKB participants participating in the AUDIT module, variants associated with increased EA were associated with decreased alcohol intake on a typical drinking day for both females and males, again, with no significant difference between sexes (overlapping CIs) (AUDIT question 2: female ß_IVW_=−0.173, 95% CI, −0.269--0.077, *P*_IVW_=4.02×10^−4^; male: ß_IVW_=−0.251, 95% CI, −0.387--0.116, *P*_IVW_=2.86×10^−4^). See Supplementary Table 6B.

Genetic variants associated with increased EA were associated with increased alcohol intake frequency for both females and males, but with no significant difference between sexes, and with evidence of residual heterogeneity and horizontal pleitropy (female: ß_IVW_=0.482, 95% CI, 0.360-0.605, *P*_IVW_=1.21×10^−14^; male: ß_IVW_=0.317, 95% CI, 0.216-0.418, *P*_IVW_=6.81×10^−10^). In contrast, variants associated with increased EA were associated with the probability of drinking alcohol with meals, and the effect was greater in males than females (female: ß_IVW_=0.132, 95% CI, 0.091-0.173, *P*_IVW_=2.50×10^−10^; male: ß_IVW_=0.246, 95% CI, 0.186-0.305, *P*_IVW_=5.50×10^−16^). See Supplementary Table 6B. For the subsample of UKB participants participating in the AUDIT module, variants associated with increased EA were associated with increased alcohol intake frequency for both females and males, with no significant difference between sexes (AUDIT question 1: female: ß_IVW_, 0.225, 95% CI, 0.063-0.388, *P*_IVW_=6.64×10^−3^; male: ß_IVW_=0.143, 95% CI, 0.001-0.285, *P*_IVW_=0.048), and decreased frequency of consuming six or more units of alcohol per occasion, with attenuated significance for females, and a non-significant effect for males (AUDIT question 3: female: ß_IVW_=−0.183, 95% CI, −0.303--0.064, *P*_IVW_=.003; male: ß_IVW_=−0.131, 95% CI, −0.285-0.024, *P*_IVW_=.098). See Supplementary Table 6B.

### EFFECTS OF EDUCATIONAL ATTAINMENT ON AD

Genetic variants associated with increased EA were associated with decreased risk of AD by almost 50% per SD unit (3.61 years schooling) (OR_IVW_=0.508, 95% CI, 0.315-0.819, *P*_IVW_=5.52×10^−3^). Magnitude and direction of the causal estimates were consistent across IVW, weighted median and weighted mode MR; the MR Egger point estimate, however, was directionally opposite but non-significant. See Supplementary Table 5. None of three AUDIT questions considered symptoms of AD, *i.e.* frequency of inability to cease drinking, frequency of failure to fulfill normal expectations, and frequency of needing a morning drink, and only one of four AUDIT questions pertaining to problematic or hazardous alcohol use, *i.e.* decreased frequency of memory loss due to alcohol, were significantly associated with increased EA (question 8: ß_IVW_=−0.113, 95% CI, −0.176--0.049, *P*_IVW_=4.92×10^−4^).

#### AD and AUDIT accounting for income

Removing variants of EA associated with average household income attenuated the significance and reduced heterogeneity but did not significantly affect the magnitude nor direction of the associations (Supplementary Table 7A). Variants associated with increased EA, but not associated with household income, were associated with decreased risk of AD by still almost 50% (per unit SD=3.61 years schooling) (OR_IVW_=0.486, 95% CI, 0.241-0.980, *P*_IVW_=0.044), and no longer with decreased frequency of memory loss due to alcohol (question 8, ß_IVW_=−0.091, 95% CI, −0.191-0.009, *P*_IVW_=0.755).

#### Sex-specific AUDIT

Genetic variants associated with increased EA were associated with decreased frequency of memory loss due to alcohol drinking in both females and males, but with no significance difference between sexes (question 8, female ß_IVW_=−0.107, 95% CI, − 0.187--0.027, *P*_IVW_=0.008; male: ß_IVW_=−0.127, 95% CI, −0.199--0.055, *P*_IVW_=5.24×10^−4^) (Supplementary Table 7B).

### BIDIRECTIONAL

#### Alcohol intake

In supplementary bidirectional analyses, genetic variants associated with DPW were not associated with EA, but variants associated with increased average weekly intake of white wine plus champagne and red wine were associated with increased EA (white wine: ß_IVW_=1.021, 95% CI, 0.765-1.278, *P*_IVW_=6.66×10^−15^; red wine: ß_IVW_=0.753, 95% CI, 0.627-0.880, *P*_IVW_=1.25×10^−31^). Conversely, variants associated with beer plus cider intake but not spirits were associated with decreased EA (beer: ß_IVW_=−0.318, 95% CI, −0.473--0.163, *P*_IVW_=5.79×10^−5^ ; spirits: ß_IVW_=−0.154, 95% CI, −0.408-0.098, *P*_IVW_=0.231). Genetic variants associated with both increased alcohol intake frequency and increased frequency of drinking with meals were associated with increased EA (intake frequency: ß_IVW_=0.212, 95% CI, 0.174-0.251, *P*_IVW_=4.49×10^−27^; meals: ß_IVW_=0.953, 95% CI, 0.786-1.119, *P*_IVW_=4.28 ×10^−29^), but residual heterogeneity and horizontal pleiotropy remained after outlier correction. Genetic variants associated with increased AD and AUDIT were not significantly associated with EA. See Supplementary Table 8.

## DISCUSSION

Using large summary-level GWAS data and complementary two-sample MR methods, we show that EA has a likely causal relationship with alcohol consumption behaviors and alcohol dependence risk in individuals of European Ancestry. More specifically, higher EA reduced binge drinking (six or more units of alcohol), the amount of alcohol consumed per occasion, frequency of memory loss due to drinking, distilled spirits intake, and AD risk. EA increased the frequency of alcohol intake, whether alcohol is consumed with meals, and wine consumption. We found evidence that our results may be driven by genetic pleiotropy in only two of the eight alcohol consumption behaviors (average weekly beer plus cider intake and alcohol usually taken with meals) and significance remained after additional analysis using EA instruments with SNPs nominally associated with either cognition or income suggest that EA may be an important factor responsible for variation in alcohol use behaviors. Consistency of our results across MR methods also strengthens our inference of causality.

Educated persons generally have healthier lifestyle habits, fewer comorbidities, and live longer than their less educated counterparts^52^ and our results suggest EA is causally associated with different likelihoods of belonging to variegated alcohol consumer typologies. We found that an additional 3.61 years of education reduced the risk of alcohol dependence by approximately 50%, which is consistent with results from small community samples^53^, and the two most recent alcohol dependence GWASs finding strong inverse genetic correlations with educational attainment^27, 54^. Notably, binge drinking significantly increases the alcohol dependence risk^55^, and distilled spirits and beer consumption account for the majority of hazardous alcohol use^56^. Furthermore, compared to wine drinkers, beer and spirits drinkers are at increased risk of becoming heavy or excessive drinkers^57^, for alcohol-related problems and illicit drug use^58, 59^, and AD^57^. Our findings related to alcoholic drink preferences, when combined with our results showing increased binge drinking, memory loss due to alcohol, and a suggestive relationship with remorse after drinking, imply a pattern of alcohol consumption motivated to reduce negative emotions or becoming intoxicated^14^.

In contrast to the often-reported positive association between EA and total amount of alcohol consumption reported from observational studies^18, 60^, we found little evidence of a causal relationship. This null finding may be reconciled by the opposing influences on alcohol intake frequency and total alcohol consumed per occasion, which, while not leading to an overall change in total consumption, nonetheless significantly affect the pattern. Our null finding regarding total consumption does support similar results from Davies et al., 2018, who used the 1972 mandated increase in school-leaving age in the UK as a natural experiment instrumental variable design to investigate the causal effects of staying in school on total alcohol consumption (from individuals in the UKB sample who turned 15 in the first year before and after the schooling age increased)^52^. Davies et al. may have found a significant effect of staying in school had they included the disaggregated behavioral dimensions of alcohol consumption behaviors. Nevertheless, even if no EA-total alcohol consumption relationship exists, studies have reported that both the specific alcoholic beverage and the pattern with which it is consumed, controlling for total consumption, independently contribute to risky health behaviors^61, 62^.

Natural experiments^52, 63^, and twin studies have found that differences in EA, even after controlling for shared environmental factors, still significantly impact mortality risk^64–66^ and recent large Mendelian randomization studies have demonstrated inverse relationships between EA on smoking behaviors^35^ and coronary heart disease (CHD) risk^34^ add to the growing body of literature suggesting a causal effect of increased EA on health and mortality. Other observational studies have linked alcohol consumption patterns to health, disease, and mortality risk^67–69^. In particular, binge drinking may have dramatic short-term consequences including motor vehicle accidents, alcoholic coma, cerebral dysfunction, and violent behavior^70^, as well as long-term effects such as hypertension, stroke and other cardiovascular outcomes^71^. A recent MR study showed that smoking mediates, in part, the effect of education on cardiovascular disease^72^, and our results suggest that differences in alcohol consumption patterns may also be another mediator. Health consequences incur significant costs with binge drinking accounting for approximately 77% of the $249 billion alcohol-related costs (lost workplace productivity, health care expenses, law enforcement and criminal justice expenses, etc.) in the United States in 2010^55^.

While we do not fully understand the underlying biological mechanisms through which the instrument SNPs influence EA, they are primarily found in genomic regions regulating brain development and expressed in neural tissue. These SNPs demonstrate significant expression throughout the life course but exhibit the highest expression during development ^36^. For example, rs4500960, which was associated with reduced EA, is an intronic variant in the transcription factor protein, T-box, Brain 1 (*TBR1*), that is important for differentiation and migration of neurons during development^36^, while rs10061788 is associated with cerebral cortex and hippocampal mossy fiber morphology^36^. It is, however, important to note that interpreting these SNPs as representing ‘genes for education’ may be “overly simplistic” since EA is strongly affected by environmental factors^36^. Our results remained when using an EA instrument with SNPs nominally associated with income removed suggesting that an individual’s genetics may impact behavior development which then increases EA^73^. Conversely, genetic estimates of EA and its correlations with other complex social phenotypes using population-based samples may be susceptible to biases, such as assortative mating and dynastic effects that provide pathways alternate to direct biological effects^40^. For example, EA-associated genetic influence on parental behavior could causally affect the child’s environment^73^. Using polygenic scores for EA, Belsky et al. 2018 recently found the mothers’ EA-linked genetics actually predicted their children’s social attainment better than the child’s own EA-linked genetics suggesting an effect mediated by environmental effects^73^. While policies are not able to change children’s genes, or their inherited social status, they can provide resources^73^, and our results suggest that interventions to increase education may help improve health outcomes through changing alcohol consumption patterns.

Notably, there was evidence for some causal effects of alcohol consumption patterns on EA, and the divergent effects again demonstrate the importance of separating drinking variables. However, we failed to find evidence that total alcohol consumed, binge drinking, or AD impacts EA, which is in line with observational studies finding no, or small effects^21^, and suggests that other studies findings a negative effect^21^ may be due to confounding. Alternatively, EA may not be sensitive enough to detect changes in schooling e.g. grade point average^21^, falling behind in homework and other academic difficulties that also reported associated with heavy drinking^74^. Further, there are currently no adolescent drinking behavior GWAS, so the temporal sequence of these analyses should be considered during their interpretation. Our findings, therefore, need replication when GWAS on adolescent alcohol consumption patterns becomes available.

Exploratory sex-specific analyses revealed differences in certain aspects of the relationship between EA and alcohol consumption. For men, the relationship between their consumption of red wine, beer, and whether they drink with meals was more sensitive to changes in EA than for women. Conversely, the reduction in binge drinking with increased EA may be driven by its effect for women since its effect on men was not significant. In addition, in women the negative effect of EA on spirit consumption was more than double its effect on men. We found no differences among the AUDIT question There are noted gender gaps in alcohol use and associated outcomes due to a combination of physiological and social factors^39^. Notably, Huerta et al. 2010 found sex-specific effects of EA and academic performance on the odds of belonging to different alcohol consumption typologies (ranging from “Abstainer” to “Regular Heavy Drinker with Problems”)^75^. The absence of any association in males may be due to their inability to model binge drinking^75^; however, our results suggest otherwise. Additionally, the recent Clarke et al. 2017 total weekly alcohol GWAS found sex-specific genetic correlation differences with an r_g_= 0.1 in men and 0.33 in women^28^. Taken together, our findings suggest EA may partially account for some of these of the observed gender gaps in alcohol consumption, but not others. We should note that the only available sex-specific EA GWAS had significant overlap (≥18.9%) with the outcome datasets, so our exploratory sex-specific analysis used the same EA GWAS combining men and women. The lack of available sex-specific AD GWAS also meant we were unable to examine differences in AD risk. Notably, the sex-specific EA GWAS demonstrated nearly identical effect sizes between men and women, which support the validity of the estimates derived from the combined-sex EA GWAS, but future studies using sex-specific instruments are required.

## STRENGTHS & LIMITATIONS

We note several strengths. We have analyzed multiple alcohol-related behavioral phenotypes which support the consistency of our results. We have implemented multiple complementary MR methods (IVW, Egger, weighted median, and weighted mode MR) and diagnostics. Consistency of results across MR methods (accommodating different assumptions about genetic pleiotropy) strengthens our causal interpretation of the estimates^76^. We also used the largest publicly available GWASs for both exposure and outcome samples; large summary datasets are important for MR and other genetic analysis investigating small effect sizes^77^. We also note limitations and future directions. There is minimal sample overlap between the exposure SSGAC GWAS and the outcome PGC GWAS (AD), but there may still be individuals participating in multiple surveys, which event we cannot ascertain with available summary level GWAS statistics. Further, the GWASs cohorts are from Anglophone countries, where beer is the preferred drink^78^; therefore, applicability to other countries with different alcohol preferences may be limited. Further still, it has been reported the UKB sample is more educated, with healthier lifestyles, and fewer health problems than the UK population^79^, which may limit the generalizability to other populations. Replication of these findings using alcohol use information from different ethnicities is necessary. EA only measured years of completed schooling; determining how various aspects of education differentially impact alcohol consumption was not possible but should be a topic of future work. Finally, alcohol consumption is not stable over time^15^; however, the alcohol consumption outcomes correspond to current drinking behavior, which may have led to the misclassification of some individuals. The current drinking also impacts the temporal relationship of our bidirectional analyses since the current alcohol intake likely occurred after maximum educational attainment for most of the participants. Future GWAS that evaluate drinking behavior during adolescence, or other longitudinal studies are necessary to confirm these findings and better elucidate the impact of alcohol intake on EA.

## CONCLUSIONS

Our data show evidence of a causal relationship between EA and patterns of drinking behavior rather than overall total alcohol consumption highlighting that drinking metrics cannot necessarily be used interchangeably. Higher EA was linked with lower binge drinking, reduced total drinks on drinking days, more frequent drinking at meals and use of moderate alcohol content beverages (such as wine). Additional education significantly reduced the risk of alcohol dependence. Alcohol consumption patterns may be significant pathways or mediators in the relationship between EA and health outcomes. In conjunction with the evidence demonstrating the causal role of education on other health behaviors, our findings suggest that increasing EA may be a useful target for prevention programs against problematic alcohol use and its consequences.

## ACKNOWLEGEMENTS

This research was facilitated by the Social Science Genetic Association Consortium (SSGAC), the Substance Use Disorders Working Group of the Psychiatric Genomics Consortium (PGC-SUD) (supported by funds from NIDA and NIMH to MH109532 and, previously, with analyst support from NIAAA to U01AA008401 (COGA)), and the Medical Research Council Integrative Epidemiology Unit (MRC-IEU, University of Bristol, UK), especially the developers of the MRC-IEU UKB GWAS Pipeline. We gratefully acknowledge their contributing studies and the participants in those studies without whom this effort would not be possible. This work was supported by the National Institutes of Health (NIH) intramural funding [ZIA-AA000242 to F.W.L]; Division of Intramural Clinical and Biological Research of the National Institute on Alcohol Abuse and Alcoholism (NIAAA). TKC, AMM and MA are supported by the Wellcome Trust (Wellcome Trust Strategic Award ‘Stratifying Resilience and Depression Longitudinally’ (STRADL) Reference 104036/Z/14/Z). The authors declare no conflict of interest.

## CODE AVAILABILITY

The analysis code in R is available on request and all data displayed in the figures are available in the Supplementary Tables.

Supplementary information is available at MP’s website.

## Notes

http://www.thessgac.org/data

http://www.med.unc.edu/pgc/results-and-downloads

http://www.mrbase.org

