## Supplemental Materials for "Educational attainment impacts drinking behaviors and risk for alcohol dependence: results from a two-sample Mendelian randomization study with ~780 000 participants"

^#^Corresponding Author:

Falk W. Lohoff, M.D.

Bethesda, MD 20892-1540

**Supplementary Information**

**Supplementary Methods** **1**. Data sources

**Supplementary Methods 2**. Alcohol Use Disorder Identification Test (AUDIT) details

**Supplementary Methods 3**. Bidirectional analysis

**Supplementary Methods 4**. Statistical analysis

**Supplementary Table 1.** Genetic correlation results from LD score regression (LDSR) of educational attainment and 20 alcohol use and dependence traits

**Supplementary Table 2.** Summary statistics for genetic instruments for educational attainment and corresponding statistics for alcohol use and dependence outcomes (main analysis)

**Supplementary Table 3.** Genetic instruments selections: educational attainment (years of schooling) accounting for average before tax household income

**Supplementary Table 4.** Description of PHESANT generated AUDIT phenotypes

**Supplementary Table 5.** Mendelian randomization results of educational attainment on alcohol use and dependence (main analysis)

**Supplementary Table 6A.** Mendelian randomization results of educational attainment on alcohol use accounting for income

**Supplementary Table 6B.** Sex-specific Mendelian randomization results of educational attainment on alcohol use

**Supplementary Table 7A** Mendelian randomization results of educational attainment on alcohol dependence and AUDIT accounting for income

**Supplementary Table 7B.** Sex-specific Mendelian randomization results of educational attainment on alcohol dependence (AUDIT)

**Supplementary Table 8.** Bidirectional analysis: Mendelian randomization results of alcohol use and dependence on educational attainment

**Supplementary Table 9.** Single SNP and leave-one-out results (main analysis)

### SUPPLEMENTARY METHODS 1. DATA SOURCES

*Education GWAS: Social Sciences Genetic Association Consortium (SSGAC)*

Full details of the methods applied for phenotype characterization, genotyping and imputation, quality control, and GWAS methods can be found in the SSGAC GWAS ^1^. All association summary statistics used in the present study are from the discovery meta-analysis (excluding the UKB replication cohorts) of the educational attainment (years of schooling, denoted *EduYears* in the GWAS) phenotype in 64 samples (combined sample size N=293 723). Of the 64, five were from single-sex cohorts, and 59 contained pooled results from mixed-sex cohorts. Only participants satisfying the following criteria were eligible for inclusion: (1) educational attainment was measured when the participant was 30 years of age or older; (2) the participant was of European ancestry, and the subject’s mother tongue was the same as the main language in the country of the cohort; (3) the participant passed the cohort’s standard quality controls, including removal of participant with poor genotyping rates or who were genetic outliers (so as to mitigate stratification concerns); and (4) all relevant covariates were available for the subject. Association analyses were conducted using linear regression; covariates included a vector of the first ten principal components of the variance-covariance matrix of the genotypic data, estimated after the removal of genetic outliers; a vector of standardized controls, including a third-order polynomial in age, an indicator for being female, and their interactions; and a vector of study-specific controls, *e.g.* major events such as wars or policy changes that may have affected access to education in their specific sample.

*Alcohol Dependence GWAS: Psychiatric Genomics Consortium*

Full details of the methods applied for phenotype characterization, genotyping and imputation, quality control, and GWAS methods can be found in the SSGAC GWAS ^2^. Individual genotypic data was collated from 14 case/control studies and nine family-based studies and summary statistics from GWAS of alcohol dependence (AD) from five additional cohorts, none of which overlapped with the SSGAC *EduYears* cohorts, nor UKB cohort. AD was defined as meeting criteria for a DSM-IV (or DSM-IIIR in one instance) diagnosis of AD. Except for three cohorts with population-based controls (N =7 015), all controls were screened for AD. Participants with no history of drinking alcohol and those meeting criteria for DSM-IV alcohol abuse were additionally excluded as controls. After initial sample and variant QC, PCA was used to identify population outliers for exclusion and to stratify European (EU) and African (AA) ancestry samples in each study. Samples were filtered for cryptic relatedness within and between cohorts. Each cohort was imputed using the 1000 Genomes Phase 3 reference panel. Imputed SNPs were then filtered for INFO score > 0.8 and allele frequency > 0.01 prior to analysis. A GWAS for AD status was performed within each ancestry stratum of each sample using an association model appropriate for the study design: for case/control studies, association analyses were performed using logistic regression with imputed dosages. Covariates included sex and principal components to control for population structure. In addition, subsets of genetically unrelated individuals were selected from each family-based cohort (*i.e.* taking one individual per family) and used to perform a conventional case/control GWAS using logistic regression. The inverse-variance weighted meta-analysis of genetically unrelated individuals was conducted in genotyped cohorts to estimate within-ancestry effect sizes for EU (N = 28 757) and AA (N = 5 799). The EU GWAS was used in the present study (N cases = 8 485, N controls = 20 272).

*Alcohol behaviors and alcohol related diagnoses: MRC-IEU UK Biobank GWAS Pipeline*

Full details of the methods applied for phenotype characterization, genotyping and imputation, quality control, and GWAS methods can be found at ^3, 4^. UK Biobank is a population-based survey of approximately 500 000 people, aged between 38 and 73 years, recruited between the years 2006 and 2010 from across the UK, with the aim of identifying the causes of human diseases in middle-age (data available at [www.ukbiobank.ac.uk](http://www.ukbiobank.ac.uk)). UK Biobank received ethical approval from the Research Ethics Committee (REC reference for UK Biobank is 11/NW/0382). The full data release contains the cohort of successfully genotyped samples (N = 488 377). Individuals with sex-mismatch (derived by comparing genetic sex and reported sex) or individuals with sex-chromosome aneuploidy were excluded (N = 814). The sample was restricted further to individuals of European ancestry as defined by an in-house k-means cluster analysis performed using the first 4 principal components provided by UK Biobank. The GWAS analysis includes the largest cluster from this cluster analysis (N = 464 708). Genome-wide association analysis (GWAS) was conducted using a linear mixed model (LMM) association method. To model population structure in the sample, 143 006 directly genotyped SNPs, obtained after filtering on MAF > 0.01, were used; genotype array and sex were adjusted for in the model.

Quality control filtering was conducted by R.Mitchell, G.Hemani, T.Dudding, L.Paternoster as described in the published protocol (doi:10.5523/bris.3074krb6t2frj29yh2b03x3wxj) ^4^. The MRC IEU UK Biobank GWAS pipeline was developed by B.Elsworth, R.Mitchell, C.Raistrick, L.Paternoster, G.Hemani, T.Gaunt (doi: 10.5523/bris.2fahpksont1zi26xosymamqo8rr) ^3^.

*Alcohol consumption (drinks per week) in the UKB: SSGAC UKB*

Full details of the methods applied for phenotype characterization, genotyping and imputation, quality control, and GWAS methods can be found in the SSGAC GWAS ^5^. All association statistics used herein are from the meta-analysis of the drinks per week construct in the UKB (sample size N = 413 343). Only participants satisfying the following criteria were eligible for inclusion: (1) the participant was of European ancestry; (2) the participant passed the standard quality controls described in the Supplementary Materials accompanying publication of the GWAS; and (3) all relevant covariates were available for the participant. The GWAS was limited to 22 autosomes. Association analyses were conducted using linear regression; covariates included a vector of principal components of the genetic relatedness matrix after application of the pre-imputation filters described above; a vector of other control variables, including controls for sex and birth year, and sex-specific birth year fixed effects; and a vector containing cohort-specific controls and technical covariates (such as dummy variables for genotyping array and genotyping batches).

*Alcohol Use Disorders Identification Test (AUDIT) in the UKB: Neale Lab UKB GWAS Round 2*

Full details of the methods applied for phenotype characterization, genotyping and imputation, quality control, and GWAS methods can be found at <http://www.nealelab.is/uk-biobank> (round 2 results, released 1 August 2018) (*see* *also* <http://www.nealelab.is/blog/2017/9/11/details-and-considerations-of-the-uk-biobank-gwas>). Starting from the 487 409 individuals with phased and imputed genotype data, the sample was filtered down to 337 199 (194 174 females and 167 020 males) QC positive individuals. Filters included removing individuals not of white British genetic ancestry; and removing closely related individuals (or at least one of a related pair of individuals), individuals with sex chromosome aneuploidies, and individuals who had withdrawn consent from the UK Biobank study. Approximately 40 autosomal million SNPs imputed from the Haplotype Reference Consortium were eligible for analysis; further restrictions to SNPs with minor allele frequency (MAF) > 0.1% and HWE p-value > 1e-10 in the 337 199 QC positive individuals, an INFO score > 0.8 (directly from UK Biobank), left 10.8 million SNPs for analysis. Phenotypes were generated using the PHEnome Scan ANalysis Tool (or PHESANT: <https://github.com/MRCIEU/PHESANT>) ^6^. Association for all phenotypes used a least-squares linear model predicting the phenotype with an additive genotype coding (0, 1, or 2 copies of the minor allele); covariates included a vector of the first 20 principal components from the UK Biobank sample QC file, and also age, age squared, inferred sex, and sex-age and sex-age squared interactions.

**SUPPLEMENTARY METHODS 2. ALCOHOL USE DISORDERS IDENTIFICATION TEST AUDIT) PHENOTYPES DETAIL**

As part of a UKB online follow up survey in 2017, 157 366 individuals responded to a mental health questionnaire, part of which included the AUDIT questionnaire. The ten-item AUDIT is as follows: (1) frequency of alcohol intake (different from alcohol intake frequency question referenced above) *i.e.* never, monthly or less, 2-4 times a month, 2-3 times a week, 4 or more times a week (completed N=117 914; UKB d-f 20414); (2) amount of alcohol drunk on a typical drinking day, *i.e.* 1-2, 3-4, 5-6, 7-9, and 10 or more (completed N=108 256; UKB d-f 20403); (3) frequency of consuming 6 or more units of alcohol, *i.e.* never, less than monthly, monthly, weekly, and daily or almost daily (completed N=108 485; UKB d-f 20416); (4) frequency of inability to cease drinking in the last year, *i.e*. never, less than monthly, monthly, weekly, and daily or almost daily (completed N=64 973; UKB d-f 20413); (5) frequency of failure to fulfill normal expectations due to drinking alcohol in the last year, *i.e*. never, less than monthly, monthly, weekly, and daily or almost daily (completed N=65 054; UKB d-f 20407); (6) frequency of needing a morning drink of alcohol after a heavy drinking session in the last year, *i.e*. never, less than monthly, monthly, weekly, and daily or almost daily (completed N=65 099; UKB d-f 20412); (7) frequency of feeling guilt or remorse after drinking alcohol in the last year, *i.e*. never, less than monthly, monthly, weekly, and daily or almost daily (completed N=65 009; UKB d-f 20409); (8) frequency of memory loss due to drinking alcohol in the last year, *i.e*. never, less than monthly, monthly, weekly, and daily or almost daily (completed N=65 029; UKB d-f 20408); (9) ever been injured or injured someone else through drinking alcohol, i.e. yes or no (completed N=118 002; UKB d-f 20411); and (10) ever had a known person concerned about or recommend reduction of alcohol consumption, i.e. yes or no (completed N=117 880; UKB d-f 20405). Summary statistics of the sample are described in Supplementary Table 4.

**SUPPLEMENTARY METHODS 3. BIDIRECTIONAL ANALYSIS DETAILS**

We extracted instruments for each of the alcohol use and dependence phenotypes from their respective GWASs, herein described, significant at genome wide significance (*P* < 5x10^-8^), pruned at the level of LD r^2^ = .001, 10000 kb clumping distance, for two-sample MR, with the following results: alcohol intake frequency (source: MRC-IEU), 99; average weekly intake of spirits (source: MRC-IEU), 4; average weekly intake of champagne or white wine (source: MRC-IEU), 4; average weekly intake of red wine source: MRC-IEU), 19; average weekly intake of beer (source: MRC-IEU), 22; drinks per week (source: SSGAC), 96; alcohol dependence (source: PGC), 3. All SNPs were present in the educational attainment outcome GWAS, excepting alcohol intake frequency, 3 not found (nor proxies). 0-2 SNPs were removed in harmonization of exposure and outcome associations. For the AUDIT phenotypes (source: Neale Lab UKB GWAS), 1-17 instruments remained after extractions of instruments, extraction of outcomes, and harmonization. Details, including single SNP and leave-one-out analyses, are available from authors. Bidirectional two-sample MR results are presented in Supplementary Table 9.

### SUPPLEMENTARY METHODS 4. STATISTICAL ANALYSIS

We used four complementary methods – inverse variance-weighted (IVW) MR, MR Egger, weighted median, and weighted mode MR – to determine the causal effects of education years on the risk of alcohol use behaviors and alcohol use disorders and discern sensitivity to different patterns of violations of IV assumptions. We refer to IVW as the main results: in the absence of pleiotropy and assuming the instruments are valid, IVW estimates are the best unbiased estimates ^8^. Consistency of results across these methods (each making different assumptions about pleiotropy) strengthens causal inference; significant divergent results may indicate bias from genetic pleiotropy.

IVW MR implements a weighted regression of the exposure SNP effects against the outcome SNP effects (*i.e*. meta-analyses SNP-specific Wald estimates from exposure and outcome GWASs), with the regression constrained to pass through the origin, and the weights derived from the inverse of the variance of the outcome effects. We used multiplicative random effects IVW MR, allowing SNPs to have different mean effects, allowing for heterogeneity due, *e.g*., to horizontal pleiotropy. Unbiased estimates of a causal effect are returned so long as horizontal pleiotropy is balanced ^8, 9^.

MR Egger extends IVW MR by not constraining the intercept to pass through the origin, allowing the net-horizontal pleiotropic effect across all SNPs to be unbalanced or directional (*i.e.* some SNPs could be acting on the outcome through a pathway other than through the exposure) ^8, 10^. MR Egger thus relaxes the assumption of “no horizontal pleiotropy” (IV assumption 2, introduction, above) and assumes only that the horizontal pleiotropic effects are not correlated with the SNP-exposure effects. MR Egger returns unbiased causal effect estimates even if the assumption is violated for all SNPs, but causal effect estimates in MR Egger are less precise (wider confidence intervals expected) than those in IVW MR. The MR Egger intercept provides an estimate of the directional pleiotropic effect ^8^.

Weighted median MR uses the median effect of all available SNPs, so that only half the SNPs need to be valid instruments (*i.e.* no horizontal pleiotropy, no association with confounders, and robust association with the exposure) to return an unbiased causal effect estimate. Stronger SNPs contribute more towards the causal estimate, with the contribution of each SNP weighted by the inverse variance of its association with the outcome ^11^. Weighted mode MR clusters the SNPs into groups based on similarity of causal effects and returns the causal effect estimate based on the cluster that has the largest number of SNPs. Weighted mode-based MR returns unbiased causal effects so long as the SNPs within the largest cluster are valid instruments. Weighted mode MR weights each SNP’s contribution to the clustering by the inverse variance of its outcome effect. Assuming the most common causal effect is consistent, the estimated causal effects would be unbiased even if all other instruments are invalid ^12^.

*Sensitivity analyses and diagnostics*

To evaluate heterogeneity in genetic instruments effects, indicating potential violations of the IV assumptions, we used the MR Egger intercept test ^13^, the Cochran Q heterogeneity test ^14^, and the MR pleiotropy residual sum and outlier (MR-PRESSO) test ^15^

MR-Egger regression provides a test for average pleiotropy: the MR Egger regression intercept can be interpreted as the average pleiotropic effect across all IVs. As pleiotropy can induce heterogeneity of the individual ratio estimates, the Cochran Q test used to identify outliers has been applied in the context of MR to detect average pleiotropy ^15^. MR-PRESSO aims to detect pleiotropic bias in MR caused by violation of the exclusion restriction IV assumption. MR-PRESSO extends the principal of the Q test, and provides a global test to detect pleiotropic bias and identify the source of the bias ^15^. Thus, MR-PRESSO identifies potential bias from horizontal pleiotropy. We used MR-PRESSO to identify the source, and then we removed the outlier variants, reran Mendelian randomization, and retested to determine whether outlier removal resolved the detected heterogeneity. We present the outlier adjusted unbiased causal estimates in Figures 2 and 3 and full MR results with outlier correction in Supplementary Tables 5-9.

**References.**

1.	Okbay A, Beauchamp JP, Fontana MA, Lee JJ, Pers TH, Rietveld CA *et al.* Genome-wide association study identifies 74 loci associated with educational attainment. *Nature* 2016; **533**(7604)**:** 539-542.

2.	Walters RK, Polimanti R, Johnson EC, McClintick JN, Adams MJ, Adkins AE *et al.* Transancestral GWAS of alcohol dependence reveals common genetic underpinnings with psychiatric disorders. *Nat Neurosci* 2018; **21**(12)**:** 1656-+.

3.	Elsworth B, Mitchell, R, Raistrick, CA, Paternoster, L, Hemani, G, Gaunt, TR MRC IEU UK Biobank GWAS pipeline version 1. University of Bristol 2017.

4.	UK Biobank Genetic Data: MRC-IEU Quality Control, Version 1. <https://research-information.bristol.ac.uk/en/datasets/uk-biobank-genetic-data-mrcieu-quality-control-version-1(fe1d820a-0114-4400-8eb1-5696f52a0b13).html>, 2017, Accessed Date Accessed 2017 Accessed.

5.	Karlsson Linnér R, Biroli P, Kong E, Meddens SFW, Wedow R, Fontana MA *et al.* Genome-wide association analyses of risk tolerance and risky behaviors in over 1 million individuals identify hundreds of loci and shared genetic influences. *Nat Genet* 2019; **51**(2)**:** 245-257.

6.	Millard LAC, Davies NM, Gaunt TR, Davey Smith G, Tilling K. Software Application Profile: PHESANT: a tool for performing automated phenome scans in UK Biobank. *Int J Epidemiol* 2017.
